# *In vivo* functional classification of PTEN variants reveals context-dependent oncogenicity and interindividual variability

**DOI:** 10.64898/2026.06.04.729913

**Authors:** David Aristizábal-Corrales, Andrés Velázquez-Mudarra, Matthias Eder, Franziska Kiem, Saúl Tamayo, Francisco J. Romero-Expósito, Mariona Cots, Alex Lugli, Alba Olaso-Llorca, Marcos Francisco Pérez, Teresa Fernández-Acero, María Isabel Rodríguez-Escudero, Caroline E. Nunes-Xavier, María Olmedo, Víctor J. Cid, Rafael Pulido, Nicholas Stroustrup, Julián Cerón

## Abstract

Gene variants, secondary mutations, and stochastic individual variability complicate cancer diagnosis, prognosis, and treatments. Here, we systematically assess the functional impact of *PTEN* cancer-related missense mutations in mammalian cell lines, yeast, and *Caenorhabditis elegans*. While cell-based assays revealed alterations in lipid phosphatase activity, CRISPR-based engineering of orthologous mutations in *C. elegans* enabled classification of variants based on organismal phenotypes and transcriptional profiles, providing a rapid framework to predict oncogenic potential. We further show that secondary mutations, such as gain-of-function of *cdc-25.1*/*CDC25A*, can enhance the phenotypic impact of specific *daf-18*/*PTEN* variants, revealing context-dependent oncogenicity. Finally, single-worm transcriptomic analyses uncovered substantial interindividual variability among isogenic animals with identical *cdc-25.1 and daf-18* mutations, linking transcriptional states to divergent phenotypic outcomes. Together, our results establish *C. elegans* as a powerful in vivo platform to integrate genetic, functional, and transcriptional information for the interpretation of cancer-associated variants.

## INTRODUCTION

The sequencing of human tumors exposes the profound complexity involved in determining genetic profiles that drive malignancy. Among factors obscuring the correlation between genotype and cancer onset and development are: i) the large number of variants in cancer genes whose functional significance is uncertain; ii) secondary mutations in the genetic background that can enhance or mitigate the effects of primary variants; and iii) interindividual variability in the phenotypic outcomes arising from factors beyond the mutational landscape.

*PTEN* encodes a lipid and protein phosphatase that plays a central role in multiple signaling pathways with functions such as cell cycle regulation, maintenance of genomic stability, or metabolic regulation (Pulido, 2015; Misra *et al*., 2021). *PTEN* variants, mostly caused by nonsense and missense mutations, are associated with three different diseases: cancer-predisposing *PTEN* hamartoma tumor syndrome (PHTS), autism spectrum disorder (ASD), and cancer (Rodríguez-Escudero *et al*., 2011). PTEN mutations in PHTS occur in the germline but two thirds of these mutations have not been detected in somatic tumors. However, most of the somatic cancer PTEN mutations have also been reported in PHTS (Post *et al*., 2020). In terms of their distribution along the protein, somatic cancer mutations are concentrated at the N-terminal phosphatase domain, whereas PHTS and ASD mutations present a broader distribution across the protein (Smith *et al*., 2019; Portelli *et al*., 2021).

A major challenge in the field is that a large proportion of *PTEN* variants correspond to Variants of Uncertain Significance (VUS). In ClinVar, a database compiling gene variants associated with clinical human diseases (Landrum *et al*., 2014), approximately 45% of single-nucleotide variants in *PTEN* are classified as VUS. Such a high proportion of *PTEN* VUS is also evident in cancer mutation databases such as OncoKB (biological and clinical annotations of somatic genetic variants in cancer) (Chakravarty *et al*., 2017; Suehnholz *et al*., 2024), cBioPortal (multidimensional cancer genomic data sets) (Cerami *et al*., 2012; Gao *et al*., 2013; de Bruijn *et al*., 2023), or COSMIC (catalog of somatic mutations in human tumors) (Forbes *et al*., 2016; Sondka *et al*., 2024). Therefore, experimental systems that enable rapid and reliable functional assessment of these variants are critically needed.

Studies of *PTEN* missense variants, initially carried out in cultured human cells and yeast, commonly by introducing a plasmid expressing the mutated version of the protein in a *PTEN*-deficient background, showed how the impact of these variants can compromise protein stability, abundance, subcellular location, and function in different ways (Matreyek *et al*., 2018; Mighell, Evans-Dutson and O’Roak, 2018; Torices *et al*., 2023). Beyond cell lines and yeast, the effects of PTEN mutations have been studied by expressing missense variants in cultured primary rat neurons, *Drosophila*, and *Caenorhabditis elegans* (Post *et al*., 2020). Alternatively, single-copy transgenes harboring missense mutations in *C. elegans daf-18* (Wittes and Greenwald, 2022) or in human *PTEN* (McDiarmid *et al*., 2018) have been inserted into the *C. elegans* genome at safe harbors but without the regulatory region of the endogenous locus. Lately, using genome editing by CRISPR-Cas9, *PTEN* missense mutations at conserved amino acids have been mimicked into *C. elegans* ortholog *daf-18* in studies focused on starvation resistance (Chen *et al*., 2022) and autism spectrum disorder (ASD) variants (González-Cavazos *et al*., 2019; Wong *et al*., 2019; Ermakova *et al*., 2022).

*C. elegans daf-18* is a well-characterized ortholog of *PTEN*, with a high proportion of amino acid conservation, particularly at catalytic regions. Importantly, similarly to PTEN, DAF-18 activity inhibits the conserved phosphatidylinositol 3-kinase (PI3K)/Akt pathway (Ogg and Ruvkun, 1998; Gil *et al*., 1999), which is aberrantly activated in distinct types of tumors promoting cancer onset and development (He *et al*., 2021). These features, plus the suitability of *C. elegans* to study cancer hallmarks, including sustained proliferative signaling or deregulation of cellular metabolism (Cerón, 2023), led us to study the functional impact of cancer *PTEN* variants affecting amino acids conserved in DAF-18. In this study, we used CRISPR-Cas9 to mimic eight mutations related to cancer according to OncoKB, including known pathogenic cancer mutations and variants of uncertain significance found in human tumors. We also investigated the effect of these variants in combination with a gain-of-function mutation in the oncogene *CDC25A* and explored how interindividual variability in the transcriptome is associated with the phenotypic outcome observed in an isogenic population carrying *PTEN* and *CDC25A* mutations.

## RESULTS

### Functional assessment of cancer-related PTEN variants in mammalian cells and yeast

To evaluate the functional impact of tumor-associated *PTEN* variants, we analyzed eight missense mutations reported in OncoKB: two variants classified as oncogenic (N48K and R130Q) were included as controls, two classified as “likely oncogenic” (R173C and Y177D), and four with unknown oncogenicity according to OncoKB (D24G, H61L, C105G, and C136F) but reported as somatic mutations in multiple tumor samples in the COSMIC database (**Fig. 1A**). All eight mutations are located within the phosphatase domain, specifically in the N-terminal region spanning amino acids 1 to 185 (**Fig. 1B**).

**Figure 1.**
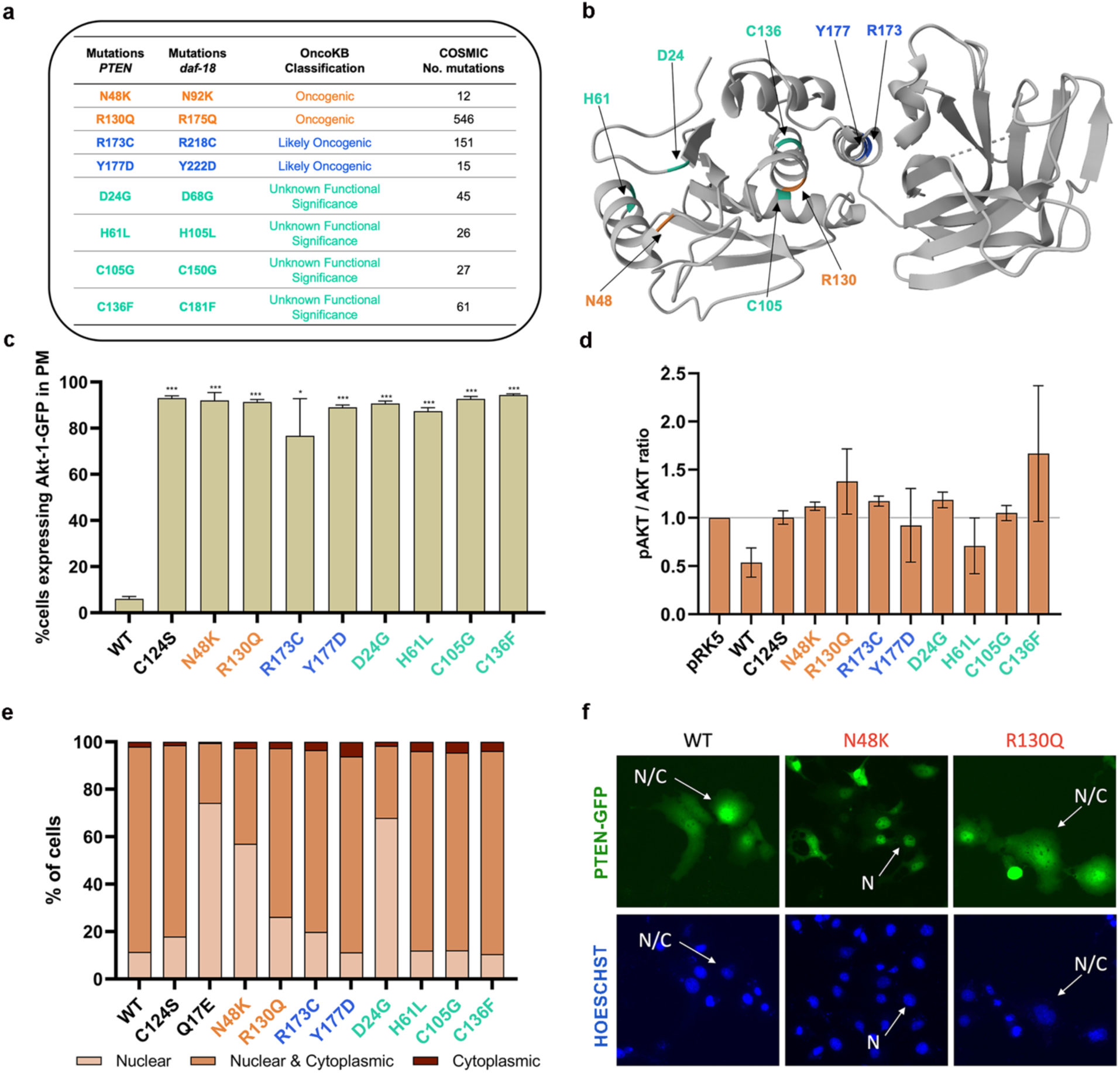
Functional assessment of cancer-related PTEN variants in yeast and mammalian cells. **a)** PTEN and *daf-18* mutations analyzed in this study, classified by predicted oncogenicity. **b)** Depiction of the 3D structure of the human PTEN protein, highlighting the localization of amino acids affected by mutations used in this study. **c)** Plasma membrane-enriched Akt1-GFP scored as an indirect output of PTEN activity in yeast cells expressing the indicated PTEN variants. Around 100 cells were analyzed per mutant. Error bars correspond to the standard deviation. T-Student tests were used for statistical significance of all *PTEN* mutants relative to WT *PTEN* (***, p<0.001; *, p<0.05) **d)** Semi-quantitative analysis of the p-AKT/AKT ratio from two Western blots was performed as an indirect measure of PTEN phosphatase activity in COS-7 cells transfected with PTEN variants. Error bars correspond to the standard deviation. **e)** COS-7 cells transfected with *PTEN-GFP* variants were analyzed by fluorescence microscopy to classify PTEN localization as nuclear, cytoplasmic, or both, according to their predominant fluorescence area. At least 50 cells per mutant were analyzed. **f)** Representative figures of PTEN::GFP localization in cells transfected with variants N48K and R130Q quantified in (e).

First, *Saccharomyces cerevisiae* was used for heterologous expression of the eight human *PTEN* variants to evaluate their impact on phosphatase activity, using the loss-of-function (LoF) variant C124S as a positive control. In this model, PTEN activity suppresses toxicity induced by human PI3K and can be monitored by the displacement of a PIP3-specific fluorescent probe (Akt1-GFP) from the plasma membrane to the cytoplasm (Rodríguez-Escudero *et al*., 2015). All missense mutants with impaired PTEN PIP3-phosphatase activity retained the Akt1-GFP signal in the plasma membrane (**Fig. 1C**), consistent with a loss-of-function phenotype. Next, we evaluated the phosphatase activity in a mammalian cell line (COS-7 cells) after transfecting the eight *PTEN* variants but this time by quantifying the abundance of phosphorylated AKT (p-AKT). PTEN lipid phosphatase activity dephosphorylates PIP3 (transforms PIP3 into PIP2), and diminished PIP3 levels reduce AKT phosphorylation. Thus, reduced PTEN activity will increase PIP3 levels and p-AKT. We observed that all *PTEN* variants expressed in cells increased the p-AKT/AKT ratio, indicating that they all affect PTEN phosphatase activity (**Fig. 1D**). Therefore, transfection of PTEN variants into unicellular models to assess phosphatase activity enabled the detection of functional alterations but did not allow discrimination between oncogenic and non-oncogenic variants as classified in OncoKB.

Finally, since PTEN has distinct functions in the cytoplasm and nucleus, we wondered if these variants would affect its subcellular location in a distinctive manner. Thus, we transfected COS-7 cells with plasmids expressing the variants fused to GFP at their C-terminus to study their subcellular distribution. We found that N48K (oncogenic) and D24G (VUS), as well as Q17E (control for nuclear localization), caused a clear increase in PTEN nuclear location. In contrast, no such increase was observed in the other variant described as oncogenic (R130Q) or the LoF variant C124S. Therefore, subcellular localization alone cannot serve as a marker of oncogenicity, although it provides valuable insight into the intracellular distribution and membrane recruitment of PTEN variants, which are key aspects of their regulatory function (**Fig. 1E**). In summary, functional assays in yeast and mammalian cells *in vitro* are informative of catalytic activity and subcellular localization of the mutated protein, but their phenotypes are not associated with the oncogenic potential of *PTEN* variants.

### Caenorhabditis elegans phenotypes of missense daf-18/PTEN cancer mutations predict their potential oncogenicity

There is a high degree of conservation at the protein level between PTEN and its *C. elegans* ortholog DAF-18. In particular, the structure of the protein tyrosine phosphatase (PTP) domain is extraordinarily similar in both proteins, supporting the functional conservation of amino acids preserved through evolution (**Fig. 2A**). Using CRISPR–Cas, we mimicked in DAF-18 the eight cancer-associated *PTEN* missense mutations located in the PTP domain selected for this study. For the sake of simplicity, the *daf-18* variants N92K and R175Q, corresponding to *PTEN* oncogenic N48K and R130Q variants, respectively, will be mentioned in this manuscript as *daf-18* oncogenic variants. Under the stereomicroscope, we observed overt RAS-related vulval phenotypes in *daf-18* mutants with these oncogenic variants (**Fig. 2B**). Worms harboring either of the two oncogenic variants, N92K and R175Q, or the deletion allele *ok480*, displayed a higher frequency of animals with defective vulva, suggesting the vulval phenotype as a good marker of *PTEN* oncogenicity for *daf-18*/*PTEN* variants. Interestingly, the variant of unknown significance C181F caused a penetrance in vulva phenotypes closer to the known oncogenic mutants, whereas the rest displayed much less severe phenotypes (**Fig. 2B**).

**Figure 2.**
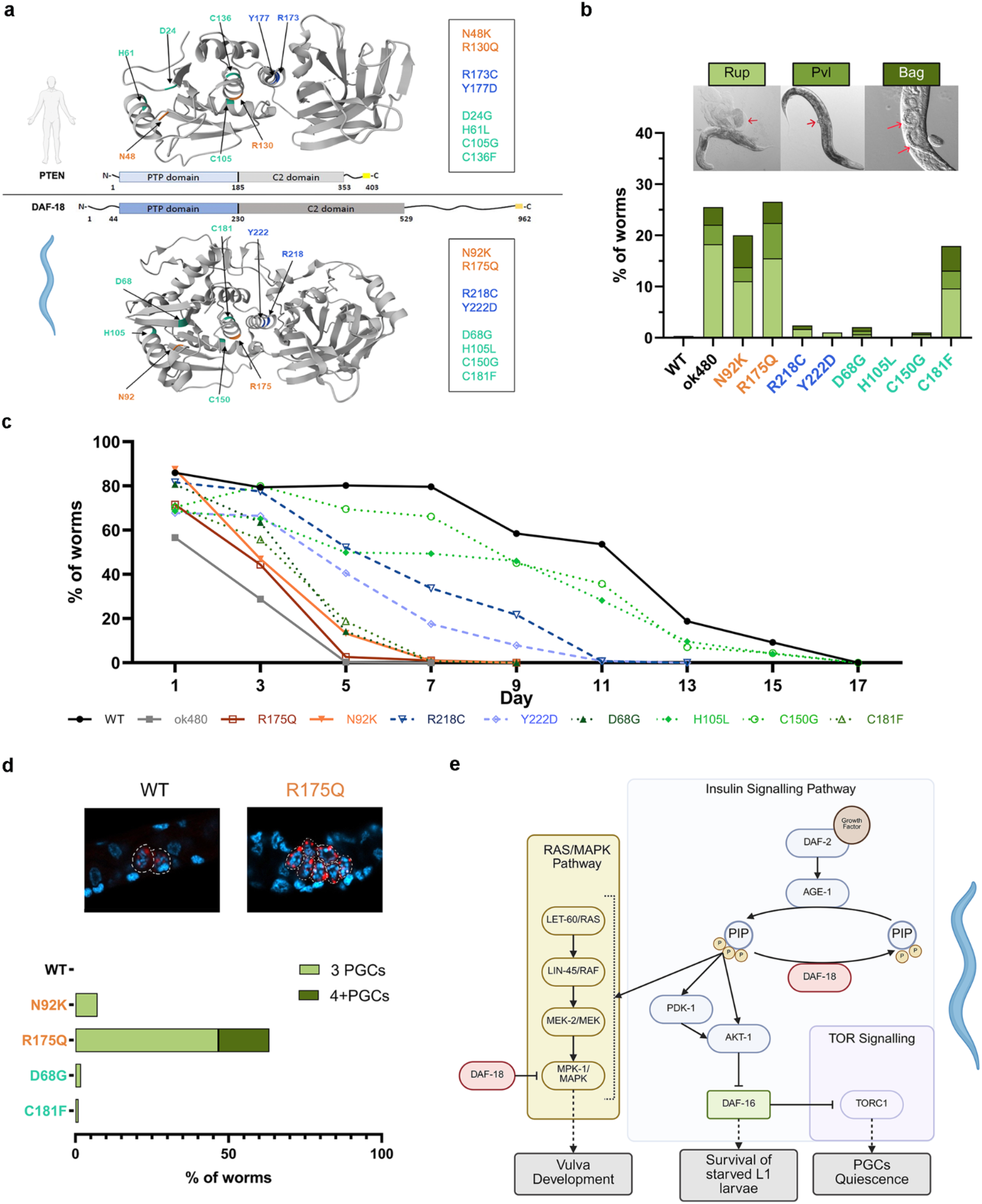
Phenotypic characterization of *daf-18/PTEN* variants. **a)** Depiction of the 3D structure of PTEN (top) and DAF-18 (bottom), and localization of amino acids targeted by mutations used in this study. **b)** Frequency of worms showing different vulva-derived phenotypes scored 4-days post-seeding L1 larvae starved for ∼24 hours. Results show the average of two independent experiments (except for Y222D; N=1). At least 50 worms were scored per variant. Representative pictures of each vulva phenotype are shown on top. Arrows indicate the phenotypic alterations: Ruptured vulva for Rup, protruding vulva for Pvl, and larvae hatched inside in the case of Bag animals. **c)** Survival after L1 starvation was measured every other day for 17 days. Results show the average of 3 independent experiments (except for ok480; N=2); n ≥70 and <200. **d)** PGCs, identified by *pgl-1*::mTagRFPT (red) expression in L1 larvae carrying the shown *daf-18* variants, were counted by using confocal microscopy after 4 days of L1 starvation arrest. Quantification shows the sum of at least two independent experiments (N≥2). At least 50 L1s were scored per variant. Nuclei were stained with DAPI (blue). **e)** Depiction of DAF-18-regulated pathways related to the phenotypes measured in our functional assays.

Since DAF-18/PTEN activity is important for the survival of L1-arrested animals in a daf-16/*FOXO* dependent manner (Chen *et al*., 2022), we wondered if a L1 starvation resistance assay would allow us to discern between our two oncogenic mutations and the six other variants. The two oncogenic *daf-18* mutations R175Q and N92K, along with the deletion mutant *daf-18(ok480)*, showed poor survival compared to the control. Two of the interrogated *daf-18/PTEN* variants of unknown functional significance (H105L, C150G) were like WT, and the likely oncogenic (R218C and Y222D) showed a mild phenotype. Remarkably, C181F and D68G mutants displayed a phenotype closer to the oncogenic variants R175Q and N92K (**Fig. 2C**).

Vulva phenotypes and L1 survival assays capture a gradient of functional effects consistent with variant oncogenicity, pointing to C136F/C181F VUS as a new potential oncogenic variant. Other PTEN activity assessments in *C. elegans* were also capable of detecting strong oncogenic variants, although not that informative for distinguishing between different levels of oncogenicity. As an example, to support the potential oncogenicity of C181F and D68G variants, we explored the proliferation of primordial germ cells (PGCs) in starved L1 larvae. PGC cell cycle is arrested during starvation, and its proliferation is controlled by the TOR signaling pathway downstream of DAF-18 (Fukuyama, Rougvie and Rothman, 2006; Fry *et al*., 2021). The known oncogenic variants, particularly R175Q, clearly produced extra PGCs during starvation, whereas only a small percentage of D68G and C181F mutants showed extra cells, and the number of PGCs in WT worms was always two (**Fig. 2D**). This is in concordance with previous studies showing that PGC proliferation is a very tightly regulated mechanism, only disturbed by strong loss-of-function *daf-18* mutants (Fry *et al*., 2021) or the double inactivation for *aak-1* and *aak-2*, which are the catalytic subunits of AMP-activated protein kinase (AMPK) (Fukuyama *et al*., 2012).

Since the survival of starved L1-stage larvae was an effective assay for inferring the oncogenicity of *daf-18/PTEN* variants, and it is a process dependent on the insulin IGF-1 receptor (InsR) orthologue *daf-2* (**Fig. 2E**), we decided to assess the capacity of *daf-18* variants to rescue the developmental delay observed in a *daf-2* mutant during the L2 stage. By using a luminescence-mediated quantitative method, previously used to measure developmental timing (Olmedo *et al*., 2015), we observed that both the deletion mutant (*ok480* allele) and the two oncogenic variants, as well as other *daf-18* variants such as D68G or H105L, and C150G to a lower extent, also rescue this *daf-2* phenotype (**Fig. S1**). Thus, most mutations at the phosphatase domain affect the DAF-18/PTEN activity responsible for rescuing the *daf-2* L2 stage delay.

In summary, across four independent assays associated with cancer-related signaling pathways in *C. elegans* (RAS, InsR, and TOR) (**Fig. 2E**), three were suitable for identifying the strong oncogenic variants, and two of these (L1 survival and vulval phenotypes) enabled discrimination among VUS daf-18 variants with distinct functional impact (**Table 1**). These results support the use of functional assays in *C. elegans* to inform on the oncogenic potential of *daf-18*/*PTEN* variants based on their capacity to phenocopy oncogenic mutations.

**Table 1.** Summary of the effect of each mutation in the four studied *C. elegans* phenotypes from most different from WT (+++++) to less (=). N.A. (Not Assessed)

| VARIANT<br>PTEN | VARIANT<br>DAF-18 | *C. elegans* PHENOTYPES |  |  |
| --- | --- | --- | --- | --- |
|  |  | L1<br>SURVIVAL | VULVA<br>DEFECTS | PGCs<br>OVERPROLIFERATION |
| N48K | N92K | ++++ | +++ | ++ |
| R130Q | R175Q | +++++ | +++++ | +++++ |
| R173C | R218C | ++ | + | N. A. |
| Y177D | Y222D | ++ | + | N. A. |
| D24G | D68G | +++ | ++ | + |
| H61L | H105L | + | = | N. A. |
| C105G | C150G | = | = | N. A. |
| C136F | C181F | ++++ | +++ | + |

### *Transcriptional profiles* of missense *daf-18*/*PTEN* mutations cluster according to oncogenicity

To further assess functional differences between variants in *C. elegans*, we performed low-input RNA-sequencing on pools of ∼45 individuals for each of the eight strains carrying the *daf-18* variants studied, along with the deletion mutant *daf-18(ok480)*, in L1 animals starved for 24 hours. We selected L1-arrested animals because they are developmentally synchronized, as they do not progress into postembryonic development in the absence of food, and DAF-18 has a role in the response to this starvation (**Fig. 2B**). After performing differential gene expression analyses (**Supplementary Table 1**), we visualized the transcriptomic profiles of the variants in a correlation heatmap (**Fig. 3A**). We found that oncogenic variants (R175Q and N92K) cluster together and next to the deletion mutant *daf-18(ok480)*. The two weakest variants in our functional assays, C150G and H105L, clustered with wild-type animals. The transcriptome of the other four mutations displaying mild phenotypes (R218C, Y222D, D68G, and C181F) in our assays clustered in pairs at intermediate positions of the heatmap. Therefore, a single amino acid change in DAF-18/PTEN variants would determine a transcriptional shift in the whole organism, and the signature of such an altered transcriptome is informative about their oncogenic potential. In the search for potential biomarkers of *daf-18*/*PTEN* variant oncogenicity in *C. elegans*, we selected genes whose normalized expression showed the strongest positive (100 genes) or negative (100 genes) correlation with phenotypic penetrance (**Supplementary Table 2**). For this analysis, we organized the variants according to the strength of their phenotype in our functional assays (**Fig. 2**), from high to low penetrance: *ok480*, R175Q, N92K, C181F, D68G, R218C, Y222D, H105L, C150G, and WT. For simplicity, we refer to this ordering as the oncogenicity rank of the variants. Among the group of genes that positively correlated with oncogenicity, we found nematode-specific genes such as *nspc-1* (the top in the list), but also several genes with human orthologues associated with cancer, according to the OncoEnrichR tool (Nakken *et al*., 2023), such as *mfb-1*/*FBX032*, *lpin-1*/*LPIN1*, or *cpr-5*/*CTSB* (**Fig. 3B** and **Fig. S2**). In the group of genes negatively correlated with oncogenicity, we identified evolutionary conserved genes related to carbohydrate metabolism, such as *gpd-3*/*GAPDH*, *aldo-1*/*ALDOC* or *enol-1*/*ENO1* (**Fig. 3B** and **Fig S2**). These genes are DAF-16/FOXO target genes that promote survival during L1 starvation (Hibshman *et al*., 2017). During L1 starvation, DAF-16/FOXO transcription factor, which is the effector of the InsR pathway, translocates to the nucleus to induce, among other things, a metabolism shift to promote survival in the absence of food (Henderson and Johnson, 2001). Thus, according to our transcriptomic data, such *daf-16* dependent remodeling of the metabolism to promote survival in L1 starved animals is affected in *daf-18* mutants, which is consistent with *daf-16* activity being impaired in *daf-18* mutants (Chen *et al*., 2022). In concordance with this hypothesis, the most oncogenic variants, *ok480*, R175Q and N92K, and in some cases C181F, showed consistent downregulation of gluconeogenesis, glyoxylate shunt and trehalose synthesis genes, and upregulation of trehalose degradation genes (*tre-5* and *tre-3*), which are transcriptional changes observed in *daf-16* mutants impairing the carbohydrate metabolism switch required for survival during L1 starvation (Hibshman *et al*., 2017) (**Fig. 3C)**. Together, these results indicate that oncogenic *daf-18*/*PTEN* variants fail to activate the DAF-16-dependent metabolic reprogramming required for starvation survival, providing a mechanistic explanation for the poor L1 survival observed in the most oncogenic variants (**Fig. 2C**).

**Figure 3.**
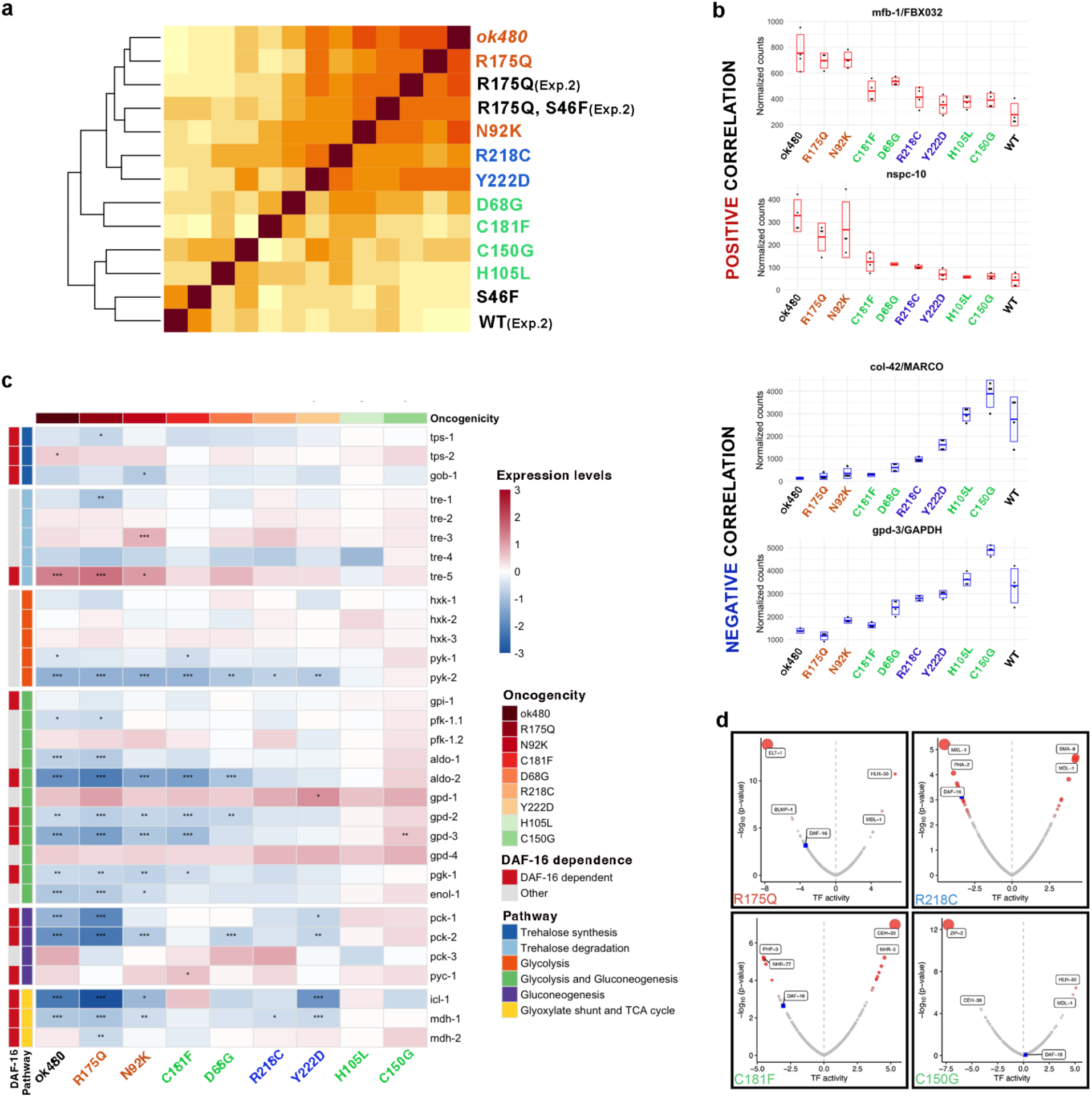
Gene expression profiles of *daf-18/PTEN* variants at L1 stage. **a)** Heatmap showing correlation clustering of the samples of different genotypes according to differential gene expressions in two different experiments. Differentially expressed transcripts were calculated with the WT sample of experiment one, as reference. Coloured text corresponds to *daf-18*/PTEN variants analyzed in the same experiment (Exp.1). Black labels correspond to a second experiment (labeled as Exp.2), in which we analyzed the transcriptomes of *cdc-25.1*[S46F], *daf-18*[R175Q], and *cdc-25.1*[S46F]; *daf-18[*R175Q] double mutants, and a second WT control. **b)** Graph shows normalized gene counts of example genes whose expression in the indicated *daf-18* mutants either positively (red boxes) or negatively (blue boxes) correlates with the oncogenicity of their corresponding *PTEN* variants. Variants are organized on the X axis from high to low oncogenicity according to our functional assays. **C)** Heatmap showing log₂ fold change in gene expression relative to wild type for carbohydrate metabolism genes identified as DAF-16-dependent targets during L1 starvation (Hibshman *et al*., 2017) across nine *daf-18*/PTEN variants in starved L1 larvae (24h). Variants are ordered from highest to lowest oncogenic potential (left to right) based on functional assays. Genes are grouped by metabolic pathway (colour-coded row annotation): trehalose synthesis, trehalose degradation, glycolysis, glycolysis and gluconeogenesis, gluconeogenesis, and glyoxylate shunt and TCA cycle. Genes previously identified as direct DAF-16 transcriptional targets are highlighted in red in the DAF-16 annotation bar. Colour scale represents log₂ fold change (blue = downregulated, red = upregulated) with values capped at ±3. Asterisks indicate statistical significance of differential expression relative to wild type (* padj < 0.05, ** padj < 0.01, *** padj < 0.001, DESeq2). **D)** Volcano plots displaying transcription factor activity scores obtained with CelEst (Perez, 2025) using gene expression data for the shown PTEN variants. On the X axis, positive values refer to positive activation while negative values correspond to lower activity. Each dot represents a transcription factor. Dot color and size reflect statistical significance: grey dots indicate low significance; red dots indicate high significance (−log₁₀(p-value)), with larger dots corresponding to higher significance. DAF-16 is highlighted using a blue square. Top 2 TFs with the highest and lowest predicted activity (mean Z-score) are labeled per genotype, along with DAF-16.

To further explore the transcriptional alterations provoked by these *PTEN* variants, we ran *Cel*EsT (Perez, 2025), a unified gene regulatory network to estimate the activity of 487 distinct transcription factors (TFs) (**Fig. 3D**). By analyzing the expression of genes regulated by these TFs, we identified transcriptional signatures for the eight variants, providing cues of altered cellular pathways. In agreement with our previous results, we observed that DAF-16 activity was strongly reduced in variants predicted to be oncogenic in our functional assays (negative TF activity values), while it remained stable in variants without detectable functional effects (**Fig. 3C and Fig. S3A**). However, when we looked at the correlation between DAF-16 TF activity and the phenotypic penetrance of the variants in our functional assays, we observed that DAF-16 activity by itself does not distinguish between strong (R175Q, N92K) and mild oncogenic variants (C181F, D68G, R218C, and Y222D) (**Fig. S3B**). This implies that other TF activities modified by oncogenic variants should be explored to study pathways implicated in PTEN inactivation and oncogenic signaling. Together, transcriptomic profiles, and expression levels in specific genes and pathways, provide an additional layer of information for classifying *PTEN* variants.

#### Impact of secondary mutations on daf-18/PTEN variants potential oncogenicity

Cancer development is caused by combinations of different mutations that function in an additive or synergistic manner to facilitate the proliferation and survival of cancer cells. Therefore, a more accurate assessment of the functional implication of variant effects in cancer onset and progress requires taking into consideration secondary mutations. As a proof of principle for this concept, we examined the effect of a *cdc-25.1*/ CDC25A mutation on *daf-18*/*PTEN* mutants. Mutations in *CDC25A* and *PTEN* co-occur in human tumor samples (according to cBioportal, p-value<0.001) (Cerami *et al*., 2012; Gao *et al*., 2013), and PTEN loss frequently co-occurs with elevated CDC25A expression (Guo *et al*., 2013; Liu *et al*., 2018), suggesting that their interaction might be relevant to cancer development.

CDC25A is a dual-specificity phosphatase that regulates the G2-to-M cell cycle transition by promoting the activation of cyclin-dependent kinases (CDKs) via dephosphorylation (Draetta and Eckstein, 1997). Therefore, CDC25A gain-of-function (GoF) in combination with PTEN tumor-suppressor loss-of-function should cooperate in promoting cell-cycle and proliferation, even in conditions of low nutrients, as in starving L1 larvae (**Fig. 4A**). The *cdc-25.1* missense variant S46F is a gain-of-function mutation (Hebeisen and Roy, 2008). Since this *cdc-25.1(cer207[S46F])* mutant causes a mild phenotype in vulva development, similar to other *daf-18*/*PTEN* variants, we investigated the consequences of combining mutations in both genes. In the case of R175Q, the strongest variant among the eight, we did not observe an enhancement of the vulva phenotype in combination with *cdc-25.1[S46F]* (**Fig. 4B**). However, a synergistic effect occurs when the *cdc-25.1* GoF mutation is combined with the mild alleles C181F and R218C (**Fig. 4B**). We did not detect any impact of *cdc-25.1[S46F]* on the WT-like variant C150G. To confirm the genetic interaction observed between *cdc-25.1[S46F]* and the VUS C181F, we quantified primordial germ cell (PGC) proliferation under starvation conditions in single and double mutants, including the oncogenic R175Q variant as a reference (**Fig. 4C**). In this assay, C181F again showed a synergistic effect with S46F, although this effect was less evident than in the vulval phenotype (**Fig. 4B**). By contrast, the double mutant R175Q; S46F did not display a synergistic effect, consistent with observations in the vulval phenotype. This likely reflects the strong functional impact of R175Q, whose phenotypic outcomes may be less susceptible to modulation by *cdc-25.1*/CDC25A gain-of-function. Consistent with this interpretation, the transcriptomes of the *daf-18___*[R175Q]; *cdc-25.1*[S46F] double mutant cluster together with *daf-18*[R175Q], whereas *cdc-25.1*[S46F] clusters closer to wild-type animals, supporting that *daf-18*[R175Q] is epistatic to *cdc-25.1*[S46F] (**Fig. 3A**). These results indicate that secondary mutations can enhance the phenotypic impact of specific *daf-18*/*PTEN* variants, allowing mild variants to cross the threshold for pathogenic classification in particular genetic contexts.

**Figure 4.**
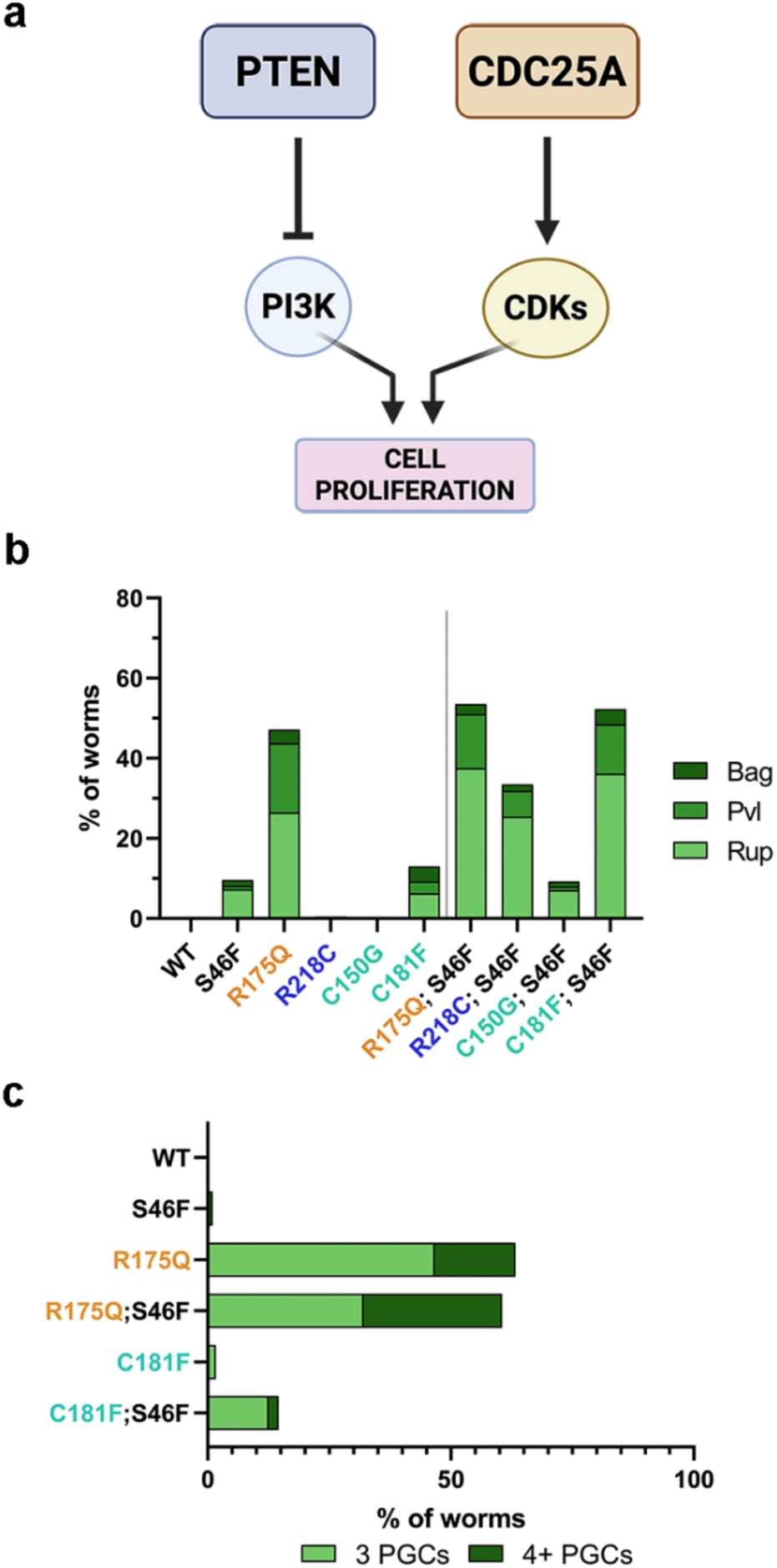
Phenotypic characterization of *daf-18* variants in the *cdc-25.1* gain-of-function context. **a)** Scheme of cell signaling interaction on cell proliferation driven by *PTEN* and *CDC25A*. **b)** Vulva related phenotypes scored in adults of the shown variant. Animals were scored 4-days post-seeding L1 larvae starved for ∼24 hours. Results show the average of three or two independent (in the case of double mutants) experiments. At least 50 worms were scored per genotype**. c)** PGCs, identified by *pgl-1*::mTagRFPT (red) expression in L1 larvae carrying the shown *daf-18* variants, were counted by using confocal microscopy after 4 days of L1 starvation. Quantification shows the average of two independent experiments. At least 30 L1s were scored per variant.

### Interindividual variability among isogenic animals

Interindividual variability represents a major challenge in cancer biology, as genetically identical tumors can display markedly different phenotypic behaviors. There is no clear genotype-phenotype association between specific *PTEN* mutations and the likelihood of developing cancer types among PHTS patients (Tan *et al*., 2012; Bubien *et al*., 2013; Nieuwenhuis *et al*., 2014). The observed variability in cancer risk among PHTS patients harboring identical PTEN mutations may arise from additional co-occurring mutations, as well as environmental or stochastic factors (Sherman *et al*., 2015; Pena-Couso *et al*., 2022). Similarly, in worms, we observed substantial interindividual variability among isogenic *cdc25.1[*S46F]; *daf-18*[R175Q] double mutant animals, which displayed a range of phenotypes, including wild-type appearance, protruding vulva (Pvl), or lethality due to vulva rupture (Rup). Taking advantage of low-input RNA-sequencing, we analyzed the transcriptome of 30 individuals for each of the three phenotypic categories: WT, Pvl, or Rup (animals with Rup phenotype were picked before they died) (**Fig. 5A**). To this end, we identified genes expressed in the germline using the WormSeq atlas ((Ghaddar *et al*., 2023); see Methods) and excluded a total of 5,827 germline-expressed genes from our dataset. This filtering was performed to remove the variation due to differences in germ cell composition of the individuals and it yielded 9,535 somatic-only expressed genes present in our RNA-seq data for downstream analysis (**Supplementary Table 3** and **Fig. S4**). By performing hierarchical clustering to study the interindividual transcriptome variability, we found better separation of the three phenotypic populations when exclusively somatically-expressed genes were considered, compared to the whole transcriptome (∼15798 genes detected). Whereas whole-transcriptome correlation heatmaps showed a narrow dynamic range (r = 0,77–1.0) with mixed phenotypic groups within each hierarchical cluster, the somatic transcriptome displayed a substantially wider correlation range (r = 0,57–1.0) and clearer phenotype-enriched clustering, with Rup individuals segregating preferentially into a distinct branch of the clustering dendrogram (**Fig. 5B** and **Fig. S5**). This demonstrates that germline filtering unmasks genuine inter-individual somatic transcriptional differences. Principal Component Analysis (PCA) using the somatic transcriptome confirmed the separation of the three phenotypic groups, with the intermediate Pvl phenotypes overlapping with both WT and Rup (**Fig. 5C**).

**Figure 5.**
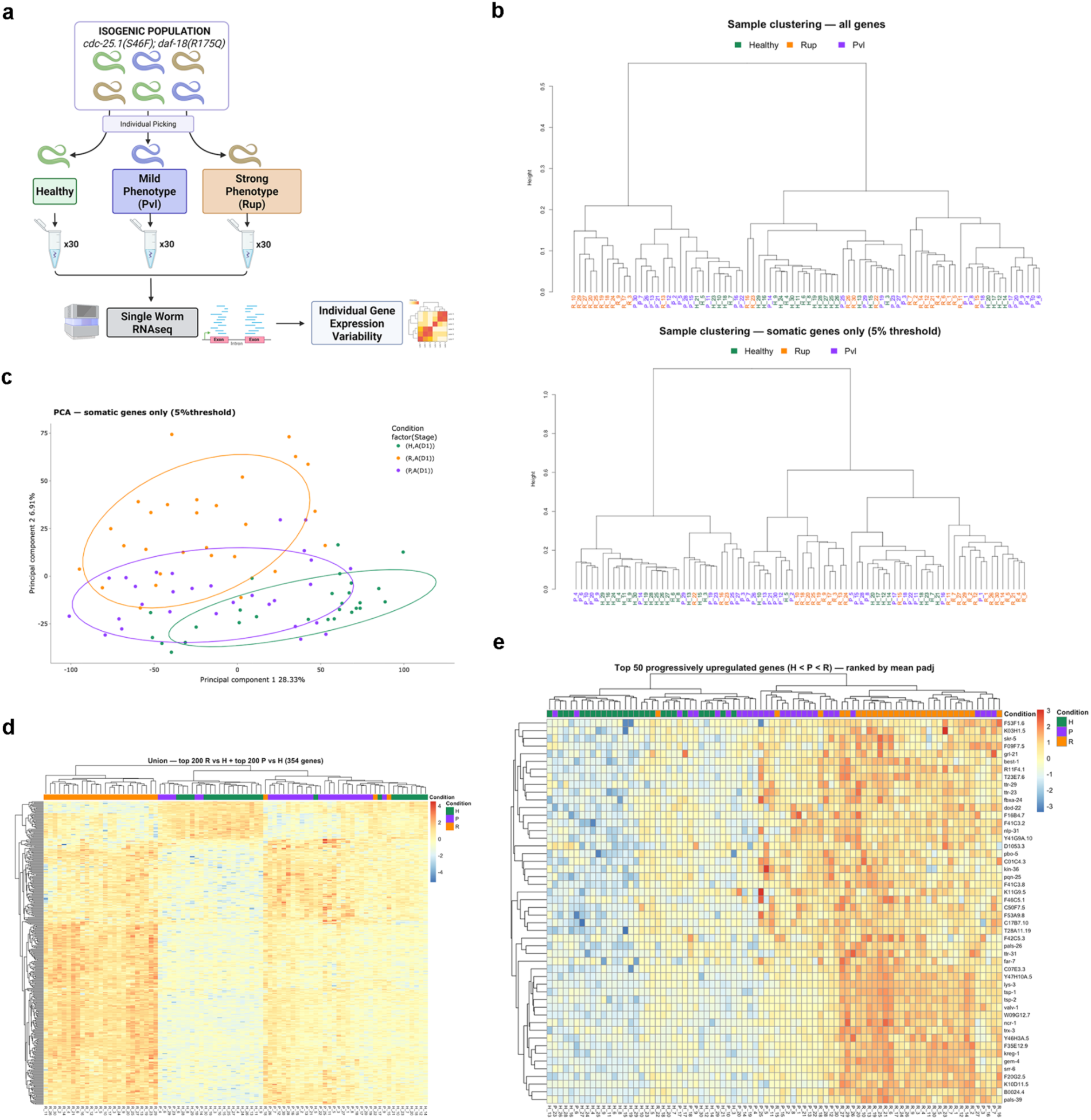
**Transcriptomic interindividual variability in *PTEN/daf-18[R175Q]*;*CDC-25*/*cdc-25.1[S46F]* double mutants**. Sample labels and dots in all panels are colored by phenotypic class: Green for Healthy, Purple for Pvl (Protruding vulva) and Orange for Rup (Ruptured vulva) animals. **a)** Schematic of the experimental design. Individual *cdc-25.1[S46F]; daf-18[R175Q*] animals were picked according to their phenotypic class — Healthy, mild phenotype (Pvl) or strong phenotype (Rup) — and subjected to single-worm low-input RNA-seq (n = 30 per class) **b)** Unsupervised hierarchical clustering dendrograms of individuals based on Pearson correlation of whole-transcriptome genes (TOP, 15.798 genes) or somatic-only gene expression (BOTTOM, 9.535 genes). Labels: H (Healthy), P (Pvl) and R (Rup). Somatic genes dendrogram clustering reveals partial phenotype-enriched grouping, the two major branches split at height ∼1.0, meaning there is a substantial transcriptional divide between the left branch (mixed H/P with a few R) and the right branch (predominantly R). Height in the Y axis = 1- r (correlation coefficient). **c)** Principal component analysis of individuals based on somatic-only gene expression. Each dot represents one individual, colored by phenotypic class (green: Healthy, red: Rup, blue: Pvl). Ellipses represent 95% confidence intervals for each group. **d)** Heatmap of 354 genes representing the topmost significantly differentially expressed somatic genes in either or both of the Pvl and Rup individuals (DESeq2, padj < 0.05 compared to Healthy). Color gradient represents expression values as z-scored log normalized counts. Columns represent individual animals ordered by hierarchical clustering; rows represent genes. Column annotations indicate phenotypic class (green: Healthy, blue: Pvl, red: Rup). **e)** Heatmap of the 50 most statistically robust progressively upregulated somatic genes, selected from the 243 genes showing a strict H < P < R mean expression pattern and ranked by average adjusted p-value across both Rup and Pvl (ranked by padj value vs Healthy). Expression values are z-scored log normalized counts. Columns represent individual animals ordered by hierarchical clustering; rows represent individual genes labeled by gene symbols. Column annotations indicate phenotypic class (green: Healthy, blue: Pvl, red: Rup)

We subsequently identified 2648 differentially expressed somatic genes (27.8% of the total somatic genes detected) in both Rup and Pvl animals when compared to the healthy-phenotype isogenic counterparts. They constitute a transcriptional signature common to both phenotypic classes, but still represent a large set of genes, reflecting the high impact that interindividual variability can have on the transcriptome of isogenic individuals (**Supplementary Table 4A**). In an attempt to identify genes associated with interindividual phenotypic differences, we extracted the top 200 most significantly differentially expressed genes (ranked by adjusted p-value) from both Rup and Pvl individuals, yielding a set of 354 genes that represents an initial core dataset associated with interindividual variability (**Fig. 5D**, **Supplementary Table 4B**). The gene expression profile of these genes highly correlates with the individual’s phenotype, as reflected in the improved clustering of the samples according to their phenotype (compare Figures 5B vs 5D). Furthermore, we found that 246 of these 354 genes showed a progressive upregulation (Healthy < Pvl < Rup), and 24 showed progressive downregulation (Healthy > Pvl > Rup) in individuals with phenotypes (**Fig. S6**), which reinforces their association with the intermediate (Pvl) and strong (Rup) phenotypic outcomes. To narrow down the list of potential markers of interindividual variability, we represented in a heatmap the top 50 genes whose expression levels increase with the penetrance of the phenotype (**Fig. 5E)**.

Thus, we identified a specific set of transcripts whose differential expression among individuals reflects the distinct phenotypic outcomes observed in animals carrying identical mutations in the cancer-related genes *daf-18* and *cdc-25.1*. Although we are unable to pinpoint the founder genes responsible for the observed variability, our results indicate that interindividual phenotypic differences within an isogenic population are reflected by pronounced variation in the transcriptome.

## DISCUSSION

A major challenge in cancer genomics is the functional interpretation of the vast number of variants identified in tumor genomes. The increasing ability to sequence tumor cells has enabled the establishment of large consortia and projects aimed at compiling data from cancer genomes (Aaltonen *et al*., 2020; Sosinsky *et al*., 2024). At the same time, this tsunami of information has provoked the development of computational tools designed to identify genomic alterations that promote tumorigenesis (Bailey *et al*., 2018; Tamborero *et al*., 2018; Martínez-Jiménez *et al*., 2020). These insights can be used for precision oncology to guide both prognosis and therapeutic strategies. Computational tools, including AI-based methods, must be trained on experimental data that support the oncogenicity of genome alterations identified in tumor genomes. Among the genome alterations promoting cancer, besides copy number variations and structural variants, genetic mutations are major cancer drivers. However, most of the genetic mutations still remain as Variants of Uncertain Significance (VUS) and an accurate re-classification as pathogenic or benign, according to ACMG/AMP standards (Richards *et al*., 2015), requires multiple independent lines of evidence. For cancer-related variants, the availability of well-validated oncogenic mutations as references facilitates the evaluation of their potential oncogenicity (Johnson *et al*., 2023).

Nowadays, CRISPR-based genome editing enables the precise reproduction of tumor-associated mutations in various cell types and, in the case of missense mutations affecting evolutionarily conserved amino acids, also across different model organisms (Serrat *et al*., 2019; Bu *et al*., 2023). The nematode *Caenorhabditis elegans* has been a relevant animal model for more than 50 years due to its amenable genetics and short life cycle. Conveniently, individuals can self-fertilize, allowing researchers to easily maintain isogenic populations (Corsi, Wightman and Chalfie, 2015). Its high homology to the human genome (Kim *et al*., 2018), particularly in genes related to human diseases, and their advantages for rapid CRISPR-Cas genome editing (Vicencio and Cerón, 2021) make *C. elegans* an attractive model for functional validation of genetic variants. *PTEN* mutations have already been mimicked in *C. elegans* by CRISPR in studies related to autism spectrum disorders (Wong *et al*., 2019; Ermakova *et al*., 2022), its protein-phosphatase activity (Chen *et al*., 2022), and the state of quiescence of the somatic gonad and germline cells in the dauer state (Tenen and Greenwald, 2019). Here, we engineered eight cancer-associated *PTEN* mutations in the *C. elegans* genome, establishing a rapid platform to assess the potential oncogenicity of *PTEN* variants while adhering to the 3Rs principles by reducing reliance on animal models. Thus, a strain mimicking *PTEN* mutation in *C. elegans*, which can be produced in two weeks, can be assessed in an L1 survival assay or scored for vulva phenotype as a robust approach to infer oncogenicity of *daf-18*/*PTEN* variants. Still, we can go deeper in exploring the effect of a *PTEN* variant in oncogenic processes by combining the *C. elegans* strain harboring the *daf-18*/*PTEN* mutation with other mutant strains. Such potential genetic interactions between variants could also be investigated by RNAi, similarly to the RNAi screen performed in mutants for *lin-35*/Retinoblastoma, a ortholog for another human tumor suppressor gene *(Ceron et al., 2007)*. These genetic interactions are relevant since they could uncover cancer vulnerabilities. In fact, CDC25 phosphatases represent a functional vulnerability in PTEN-inactivated cancers, including triple-negative breast tumors (Liu *et al*., 2018). Consistent with our results showing distinct levels of genetic interactions, tumors harboring specific PTEN variants may be more susceptible to treatment with CDC25A inhibitors than others.

Moreover, our study faces the question of interindividual variability among isogenic tumors, which is a critical and complex question in cancer prognosis. The impact of environmental conditions and the stochastic variation in the penetrance of phenotypes of isogenic animals have been studied in *C. elegans*. Several mechanisms have been proposed to explain this interindividual variability among isogenic mutant animals (Burga, Casanueva and Lehner, 2011; Burga and Lehner, 2012; Casanueva, Burga and Lehner, 2012), and also in cancer cells (Whiting *et al*., 2024), including differential compensation by other pathways, variation in chaperone expression levels, differences in stress responses, or epigenetic modifications. We studied the transcriptome of double mutants for *daf-18* and *cdc25.1* individuals displaying phenotypic heterogeneity. We found that, at the day-one adult stage, the expression of roughly one third of the somatic genes detected was deregulated in animals displaying Rup or Pvl phenotypes.

Therefore, it is challenging to identify the specific “founder” genes responsible for triggering such widespread transcriptomic changes. However, our data suggests that even after excluding germline-expressed genes (which can also be expressed in the soma but represent a major source of interindividual variability), large-scale transcriptional changes occur in isogenic nematodes during postembryonic development, which is a highly proliferative system like tumors.

Our study provides experimental evidence supporting that *PTEN* C136F and D24G, which were classified as variants of uncertain significance, have oncogenic potential since they phenocopy known *daf-18*/*PTEN* oncogenic variants, and display similarities to these variants of reference in their transcriptomic profiles. The existence of tumor suppressor genes with specific functional activities related to cancer, such as DNA repair in the case of BRCA1, facilitates the experimental validation of VUS oncogenicity using *in-cellulo* systems (Hu *et al*., 2022). However, PTEN functions in cancer are much more diverse and dependent on the context, making a multicellular organism such as *C. elegans* an advantageous model for functional studies of gene variants. Moreover, we show how the combination with secondary mutations and factors causing interindividual variability can be relevant for cells harboring genetic variants that, in principle, seem to lack any major functional impact. Therefore, we believe that the animal model *C. elegans* can extend its impact in cancer research, offering a valuable layer of experimental evidence on gene variants to feed databases and AI tools.

## MATERIALS AND METHODS

### C. elegans strains

The wild-type strain N2 and RB721 *daf-18(ok480) IV* were obtained from the Caenorhabditis Genetics Center (CGC). We used CRISPR-Cas technologies to generate eight different strains carrying the individual *daf-18/PTEN* mutants. Some of these mutants were also inserted in the *cdc-25.1(cer207[cdc-25.1[S46F]]) I* background, which we also generated in our lab. Some of them were also inserted in the *pgl-1(sam52[pgl-1::mTagRFPT::3xFlag]) IV* strain, which we obtained by crossing N2 with the DUP121 strain, obtained from (Marnik *et al*., 2019). The full list of strains can be found in **Supplementary Table 5A**.

### Assessment of the activity of PTEN variants in yeast

The YPH499 (*MATa ade2-101 trp1-63 leu2-1 ura3-52 his3-Δ200 lys2-801*) *Saccharomyces cerevisiae* strain was grown in synthetic complete (SC) medium (0.17%) yeast nitrogen base without amino acids, 0.5% ammonium sulfate, provided with the appropriate supplements to maintain auxotrophic marker-based plasmids, and 2% glucose (SD), galactose (SG), or raffinose (SR), as required. Yeast transformation was achieved by the standard lithium acetate protocol.

For GFP-AKT1 membrane localization, as a surrogate indicator of cellular PIP3, which is converted to PIP2 by catalytically active PTEN, transformant YPH499 cells bearing pYES3-Akt1 (Rodríguez-Escudero *et al*., 2011), pYES2-PTEN (WT or variants) and YCpLG myc-p110α-CAAX were grown overnight in liquid SR medium lacking Trp, Ura and Leu to maintain all three plasmids, and 2% galactose was added for 5h to induce the expression the three heterologous proteins from the GAL1 promoter. GFP-AKT1 plasma membrane localization was assessed by standard fluorescence microscopy. At least 100 cells were analyzed per experiment.

### Mammalian cells assays

For all experiments performed in mammalian cells, the COS-7 cell line was used. Cells were cultured at 37 °C with 5% CO₂ in DMEM supplemented with 5% heat-inactivated fetal bovine serum (FBS), 1 mM L-glutamine, 100 U/mL penicillin, and 0.1 mg/mL streptomycin. For the expression of PTEN variants, the pRK5 vector was used. The directed mutagenesis was achieved by inverse PCR (Mingo *et al*., 2018). The product was digested with *DpnI* (New England Bio Labs) overnight at 37°C for the following transformation with competent *E. coli* DH5⍺. The extraction and purification of these plasmids was performed with the NucleoSpin® Plasmid EasyPure (Macherey-Nagel) kit. Presence of the desired PTEN mutations was ratified by digestion with different digestive enzymes and later sequencing of the samples. The GenJet^TM^ transfection protocol was followed to transfect COS-7 cells.

To study pAKT1 levels, COS-7 cells were co-transfected with the different PTEN variants and HA-AKT1, and Western blot analyses were performed using monoclonal antibodies against phospho-Ser473-AKT, AKT (Cell Signaling Technologies, USA), and PTEN 6H2.1. As a protein loading control, GAPDH (glyceraldehyde-3-phosphate dehydrogenase) expression was analyzed. To determine the predominant subcellular localization of PTEN variants, a PTEN-GFP plasmid was modified by introducing the mutations of interest. The nuclei were identified by Hoechst staining. They were visualized by fluorescence microscopy using a LEICA DMIL LED microscope. All images were acquired with a 20X objective. To quantify the nuclear/cytoplasmic distribution of PTEN, at least 50 positive cells were analyzed in each experiment. The cells were classified by the area with enriched GFP signal (nuclear, cytoplasmic, or both).

### Mutant strains and CRISPR guides

*C. elegans* strains are maintained in NGM plates seeded with OP50 *E. coli* as a food source between 15-25°C. Strains mimicking human mutations were created by CRISPR-Cas9. The preparation of CRISPR Mixes (**Supplementary Table 5B and 5C**) involves the synthesis of two gRNAs (for the targeted gene and for the phenotypic reporter *dpy-10*), followed by the formation of RNP Complexes. Repair templates for the corresponding mutation of interest and for the reporter for selection (*dpy-10*) were also added to the mix. CRISPR Mixes were microinjected into the germline of adult worms, and the progeny were further scanned and genotyped to select for the mimicked mutations.

### L1 synchronization

Gravid adults are collected off plates with M9 buffer and washed to eliminate excess bacteria. Worm pellets diluted in 1mL of M9 buffer are treated with 5 mL of bleaching solution (1.2% bleach, 0.625M NaOH in ddH_2_O) until most of the cuticle is broken down and the embryos are released (normally less than 5 min). The embryos obtained are washed at least 3 times with M9 buffer and left to hatch overnight without food in the same buffer at 20°C.

### L1 Survival

Worm populations were bleached and left in 2mL of M9 rotating at 20°C in 15mL Falcon tubes. Every other day, across 15-17 days, between 150-200 L1 stage worms were seeded in NGM agar plates with OP50 to feed the worms (exact number of worms seeded per plate is scored). After 48 hours, the number of alive worms was scored to calculate the survival percentage.

### Analysis of postembryonic developmental timing

Developmental timing of individual worms was measured using a bioluminescence-based assay as previously described (Olmedo *et al*., 2015). To obtain age-synchronized embryos, 10–15 adult hermaphrodites were transferred to a fresh NGM plate and allowed to lay eggs for one hour. Individual embryos were then placed into wells of a 96-well plate containing 100 μl of S-basal medium (supplemented with 10 μg/ml cholesterol) and 200 μM luciferin. After all embryos were loaded, 100 μl of S-basal containing 20 g/l E. coli OP50-1 was added to each well. Plates were sealed with a gas-permeable membrane and incubated in a luminometer (Berthold Centro XS3) housed within a cooled incubator (Panasonic MIR-254).

### Vulva phenotyping

Synchronized L1 larvae were maintained in 2mL of M9 buffer at 20°C to hatch in starvation conditions for ∼24h. The starved L1s are then seeded (around 200 worms) in NGM plates with OP50. After 4 days, when they reach day-2 adulthood, the total number of worms and their vulva phenotypes (Bag, Pvl, Rup) were scored.

### Germ cell precursor scoring

The *daf-18* mutations were introduced in worms carrying the *pgl-1*::mTagRFPT::3xFLAG reporter for germ cells (Marnik *et al*., 2019). Worm populations carrying the inserted mutations were bleached, and the obtained embryos hatched in 2mL of M9 buffer rotating at 20°C for 4-days starvation. Starving L1s are then fixed by several incubations with 2% formaldehyde, PBS-Triton (0.5%), and 70% Ethanol and washes with PBS-Tween (0.1%). Fixed worms were mounted in microscopy slides with the SouthernBiotech DAPI Fluoromount-G mounting media. Precursor Germ Cells were scored using confocal fluorescence microscopy.

### RNA sample collection and processing for sequencing

Collection of L1 pools samples of the corresponding variants mentioned in the results was done as follows. Embryos were obtained by bleaching and incubated in 2mL of M9 buffer rotating at 20°C overnight without food. Hatched L1s were counted 24h later, and four PCR tubes with ∼45 worms were prepared per strain. These samples were spun down, and the worm pellets were diluted in 65μL of Lysis buffer. Samples were snap frozen in dry ice-cold ethanol and stored at −80°C until further preparation.

For the interindividual variability study, embryos from an isogenic strain (CER751) were obtained by bleaching and kept in an M9 buffer rotating overnight without food. After ∼24h, hatched L1 worms were seeded on plates with food, and when they were adults, 96h after plating, they were individually picked according to their phenotype. Ruptured worms were still alive and moving (the grade of bursting varied, but in all cases, at least 1/4 of the intestine was bursted out of the vulva); Pvl were worms displaying small protruding vulvas; Healthy fertile wild-type looking worms were mostly early adults with a single row of embryos. For each phenotype, 30 worms were picked and transferred individually to PCR tubes with 8μL of lysis buffer. Samples were snap frozen in dry ice-cold ethanol and stored at −80°C until further preparation.

For Low-Input RNA sequencing, samples were processed following the protocol described by (Serra *et al*., 2018), with the inclusion of ERCC spike-ins added to the lysis buffer at a final dilution of 1:40,000. Lysis was conducted at 65 °C for 10 minutes, followed by enzyme inactivation at 85 °C for 5 minutes. cDNA libraries were prepared using the Smart-seq2 protocol and subsequently purified using in-house–prepared SPRI paramagnetic beads (functionally equivalent to AMPure XP beads, Beckman Coulter) at a bead-to-sample ratio of 0.8. Library size distribution was evaluated using a TapeStation 4150 (Agilent), and cDNA concentrations were measured with Quant-iT (Invitrogen) on a Tecan plate reader. Sequencing libraries were then generated from the cDNA using Nextera tagmentation and PCR amplification with indexed primers, following the Nextera DNA library preparation protocol (Illumina). These libraries were purified twice using in-house SPRI beads at a bead-to-sample ratio of 0.9. Library size profiles were reassessed on a TapeStation 4150 (Agilent), and concentrations were quantified using Quant-iT (Invitrogen) on a Tecan plate reader. RNA-seq libraries were pooled in equal masses, and sequencing was carried out on an Illumina NovaSeq 6000 using an S2 100-cycle kit (Illumina), generating in average >2.0 × 10^6 2x52-bp paired-end reads per sample.

### RNAseq Analysis

RNA seq data were analyzed using R v4.4.1 (R Core Team, 2024). For both L1 pools and Single Adult samples, RNAseq reads from Illumina NovaSeq were aligned to the C. elegans WormBase reference genome (WS265 release) using STAR v2.6.0c (Dobin *et al*., 2013). Gene-level counts were then derived from these alignments with featureCounts v2.0.0 (Liao, Smyth and Shi, 2014). Raw counts for each mapped gene were normalized using the scran package v1.32.0 (Lun, McCarthy and Marioni, 2016), which computes pooling-based size factors robust to zero counts in low-input data. Genes with fewer than 5 counts in more than 50% of samples were excluded prior to normalization. Samples with total read counts below one-tenth of the median library size were excluded. Differential expression analysis was performed using DESeq2 v1.44.0 (Love, Huber and Anders, 2014), with scran-derived size factors supplied directly to replace DESeq2’s internal normalization. Dispersion was estimated using the default parametric fit, and significance was assessed using the negative binomial Wald test. All comparisons were made relative to the wild-type condition (WT, N2 strain in the case of L1 pools or Healthy individuals in the case of Single Worm seq). Results were extracted for each pairwise condition contrast and annotated with gene symbols using the *C. elegans* annotation package org.Ce.eg.db v3.19.1 (Carlson, 2024) via the Bioconductor AnnotationDbi interface. Either normalized counts or differential expression information from DESeq2 (log2 fold change and p-adjusted value) were used for the downstream analysis mentioned in the text. To identify gene lists associated with *daf-18* variant oncogenicity in L1 pools data (described on Fig 3B and Supplemental Table 2) a per transcript linear regression was run on the log normalized counts with the oncogenicity of each variant as a ranked categorical variable.

To remove the confounding effects of differential germline cell composition between individuals for the analysis of interindividual variability using single worm RNA seq data, we filtered the raw count expression matrix to retain only genes expressed exclusively in somatic tissues. Germline-expressed genes were defined as genes detected in more than 5% of germline cells in the (Ghaddar *et al*., 2023) whole-body adult C. elegans scRNA-seq atlas dataset. Using this threshold we identified 5,747 germline-expressed genes across nine germline cell type clusters: mitotic, meiotic and apoptotic germ cells, differentiated germ cells, oocytes, spermatocytes, spermatids, mature sperm and syncytial pachytene spermatocytes. To assess the robustness of the germline gene filtering strategy, PCA was performed using germline genes defined at both 5% and 10% expression thresholds. Results were consistent across both thresholds, and the 5% threshold was used for all subsequent analyses. This threshold was validated against three independent germline transcriptome datasets — (Wang, Gerstein and Snyder, 2009; Ortiz *et al*., 2014; Tzur *et al*., 2018) — capturing 75.6%, 50.1% and 66.0% of their respective germline gene catalogues (Supplementary Table 4). This filtering yielded 9,535 somatic only-expressed genes present in our RNAseq data for downstream analysis.

---

**Figure S1.**
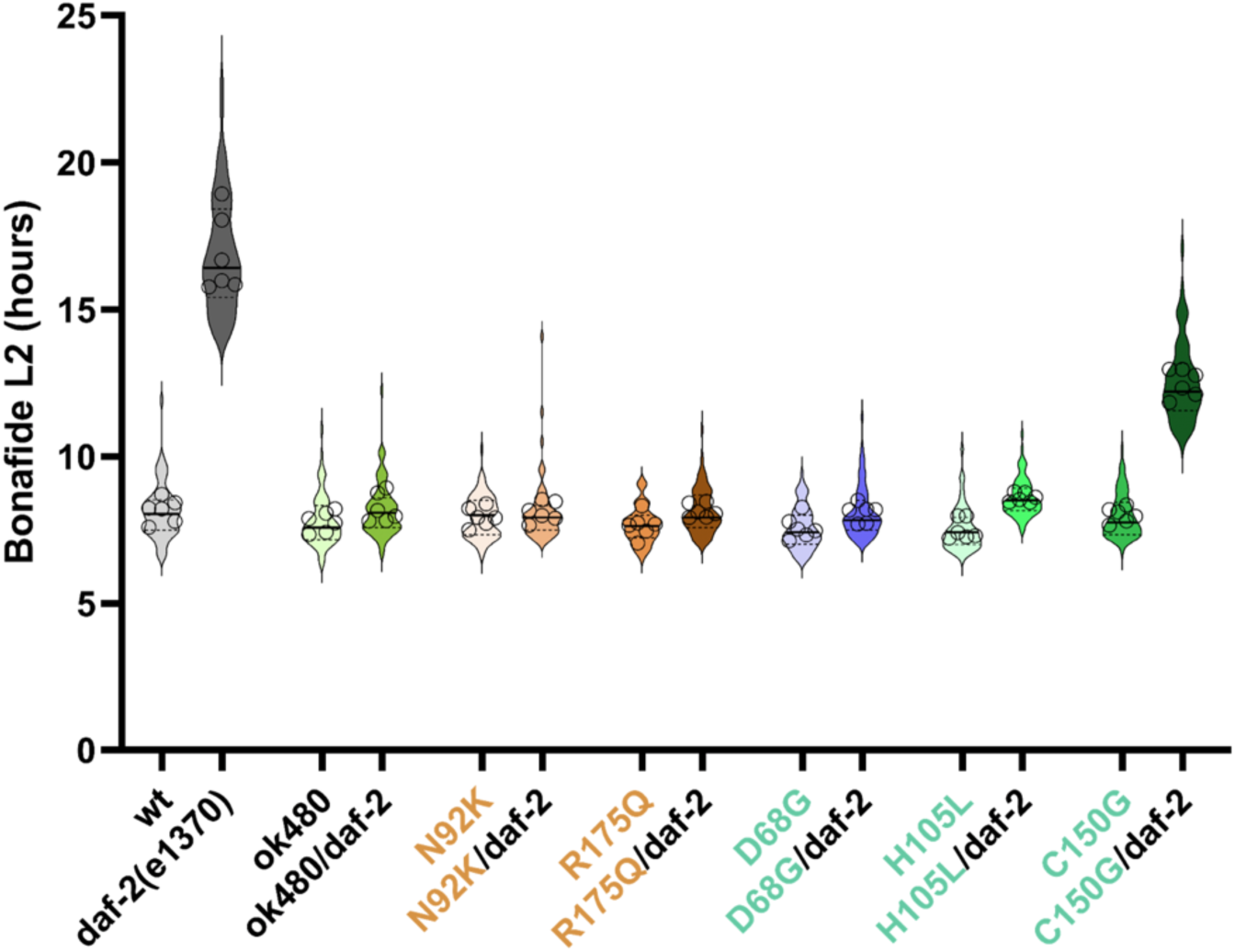
Developmental timing analysis of daf-18/PTEN variants in a daf-2 mutant background. Violin plots showing the duration of the L2 larval stage (bonafide L2, hours) for wild-type (wt), daf-2(e1370), daf-18(ok480) null allele, and six daf-18/PTEN missense variants (N92K, R175Q, D68G, H105L, C150G), each assessed alone and in combination with daf-2(e1370). Developmental timing was measured using a luciferase-based bioluminescence method as described in Olmedo et al. (2015). daf-2(e1370) single mutants displayed a significantly prolonged L2 stage compared to wild-type, consistent with the known role of DAF-2/insulin receptor signaling in developmental timing. Loss of daf-18 (ok480 allele) and oncogenic variants N92K and R175Q suppressed the daf-2 developmental delay, restoring L2 duration to near wild-type levels, consistent with their strong loss-of-function impact on DAF-18/PTEN activity. D68G showed a similar suppression, while H105L and C150G showed partial rescue. Individual data points are shown as circles overlaid on each violin. n ≥ 10 animals per condition. Horizontal lines within violins indicate median and interquartile range.

**Figure S2.**
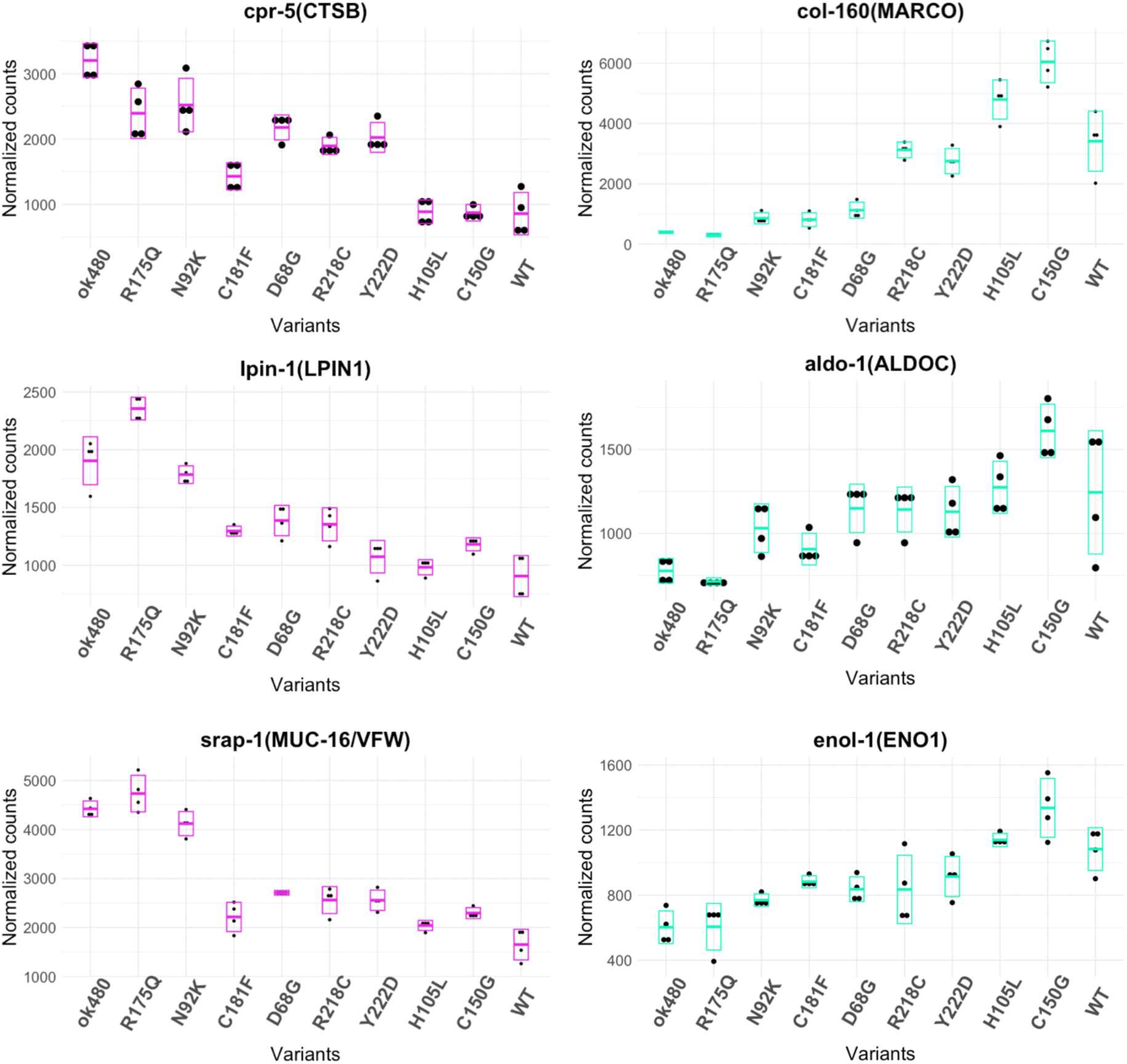
Other genes whose expression correlate with *daf-18* variant oncogenicity during L1 stage starvation. Graph shows normalized gene counts of genes whose expression in the indicated *daf-18* mutants either positively (red boxes) or negatively (blue boxes) correlates with the oncogenicity of their corresponding PTEN variants. Variants are organized in the X axis from high to low oncogenicity according to our functional assays.

**Figure S3.**
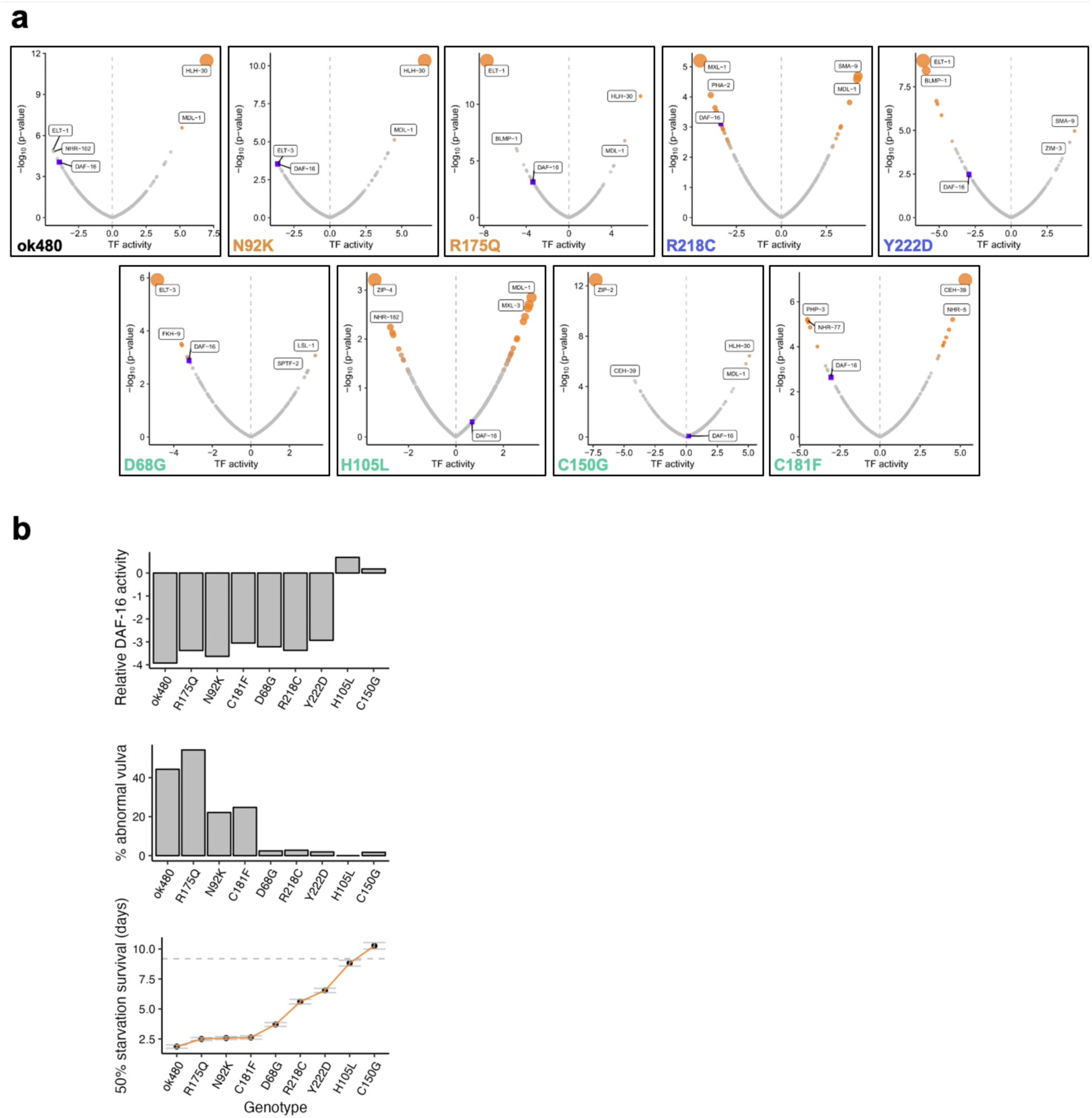
Transcription factors activities in PTEN mutants in L1 stage. **a)** Volcano plots displaying transcription factor activity scores obtained with Cel/EsT (Perez, 2025) using differential expression data for the shown PTEN variants at L1 stage. On the X axis, positive values refer to positive activation while negative values correspond to lower activity. Dots labeled in red correspond to transcription factors that show highly significant p value of activity. DAF-16 activity, a known DAF-18 target, is labeled in blue. **b)** Correlation between DAF-16 activity and the oncogenicity of *daf-18/*PTEN variants. In all plots, variants are ranked on the X axis according to their oncogenicity, from high to low. Top plot: Bars represent transcription factor activity scores for DAF-16 obtained with CelEst for each variant. Middle plot: Bars represent the sum of all vulva related phenotypes observed on each variant from Fig 2b. Bottom plot: Represent the L1 Starvation survival for each *daf-18/*PTEN variant. Y axis shows the day at which 50% of the worms are still alive during L1 starvation. Data was taken from Fig 2c.

**Figure S4.**
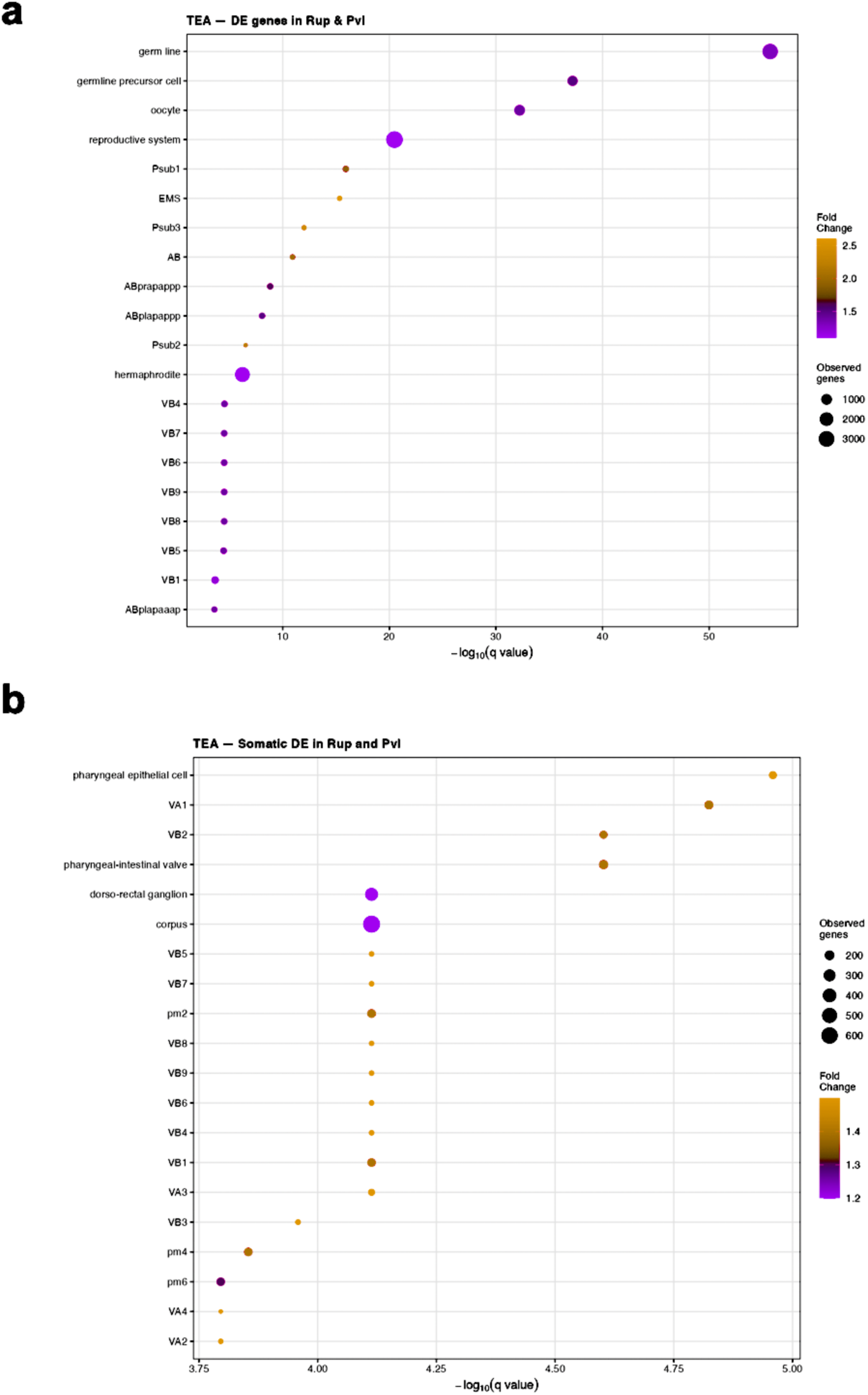
Tissue enrichment analysis (TEA) of genes commonly differentially expressed in Rup and Pvl — whole transcriptome vs somatic only genes. TEA for dysregulated differentially expressed genes comparing genes sets from **a**) whole transcriptome vs **b)** somatic only genes. For **a**) and **b**) Bubble plot showing tissue enrichment of genes significantly DE (padj < 0.05) in both Rup and Pvl. The x-axis shows −log₁₀ (q value); bubble size reflects the number of observed genes per term; color indicates enrichment fold change. In **a**) germline and reproductive tissues dominate the enrichment, reflecting the strong germline expression signal present in the unfiltered whole transcriptome. In **b**) all germline related categories are lost, confirming the somatic only filtering of our data.

**Figure S5.**
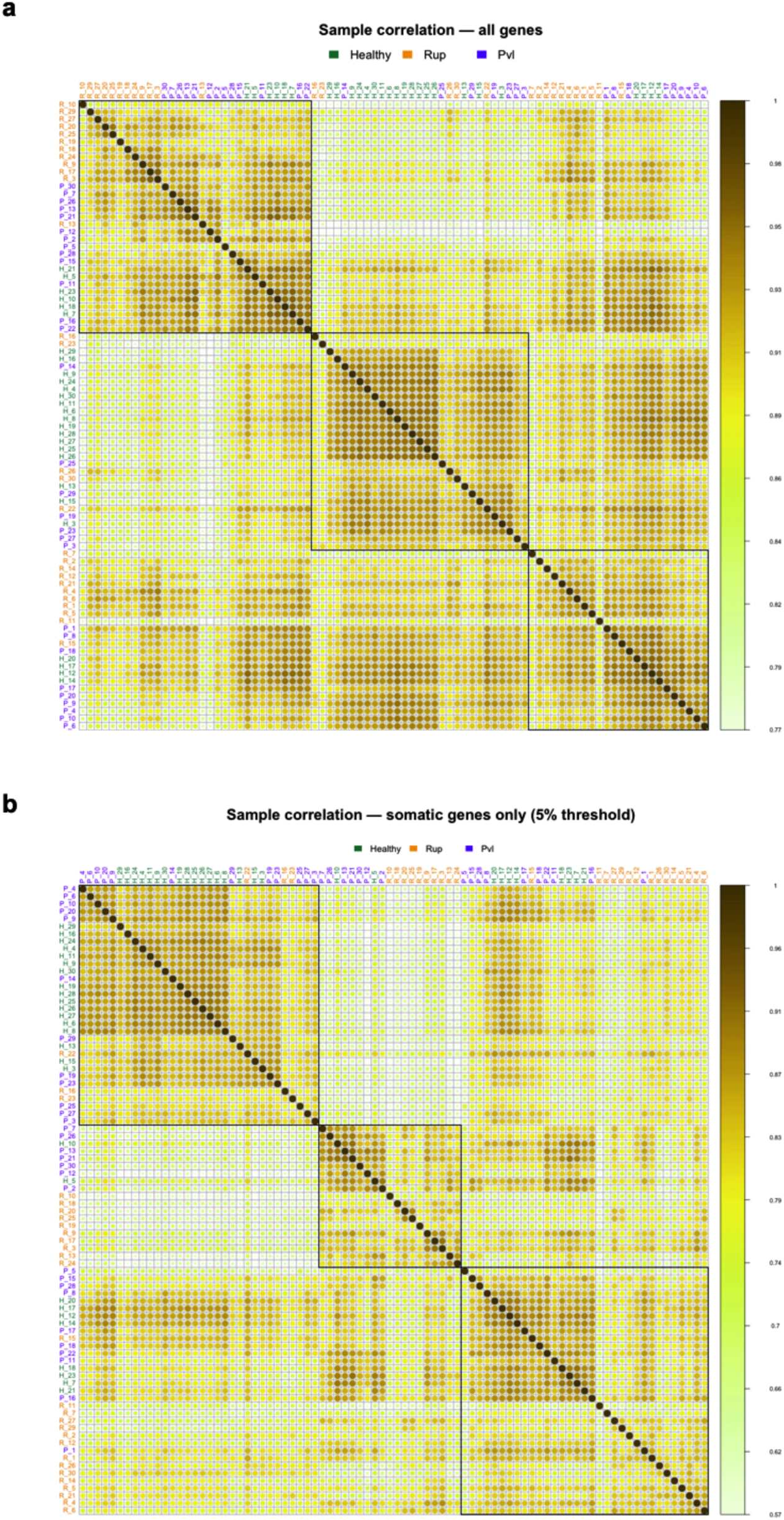
Single Worm Sample correlation heatmaps before and after germline gene filtering. Pairwise Pearson correlation matrices of all individuals based on (**a**) the whole transcriptome (∼15,000 detected genes; r = 0.77–1.0) and (**b**) somatic-only genes following removal of germline-expressed genes (9,535 genes; 5% threshold, Ghaddar et al. 2023; r = 0.57–1.0). Circle size and color intensity reflect pairwise correlation strength. Black boxes highlight major hierarchical clusters. Sample labels are colored by phenotypic class (green: Healthy, red: Rup, blue: Pvl). The wider correlation range and clearer phenotype-enriched clustering in (**b**) compared to (a) demonstrates that germline filtering unmasks genuine inter-individual somatic transcriptional differences.

**Figure S6.**
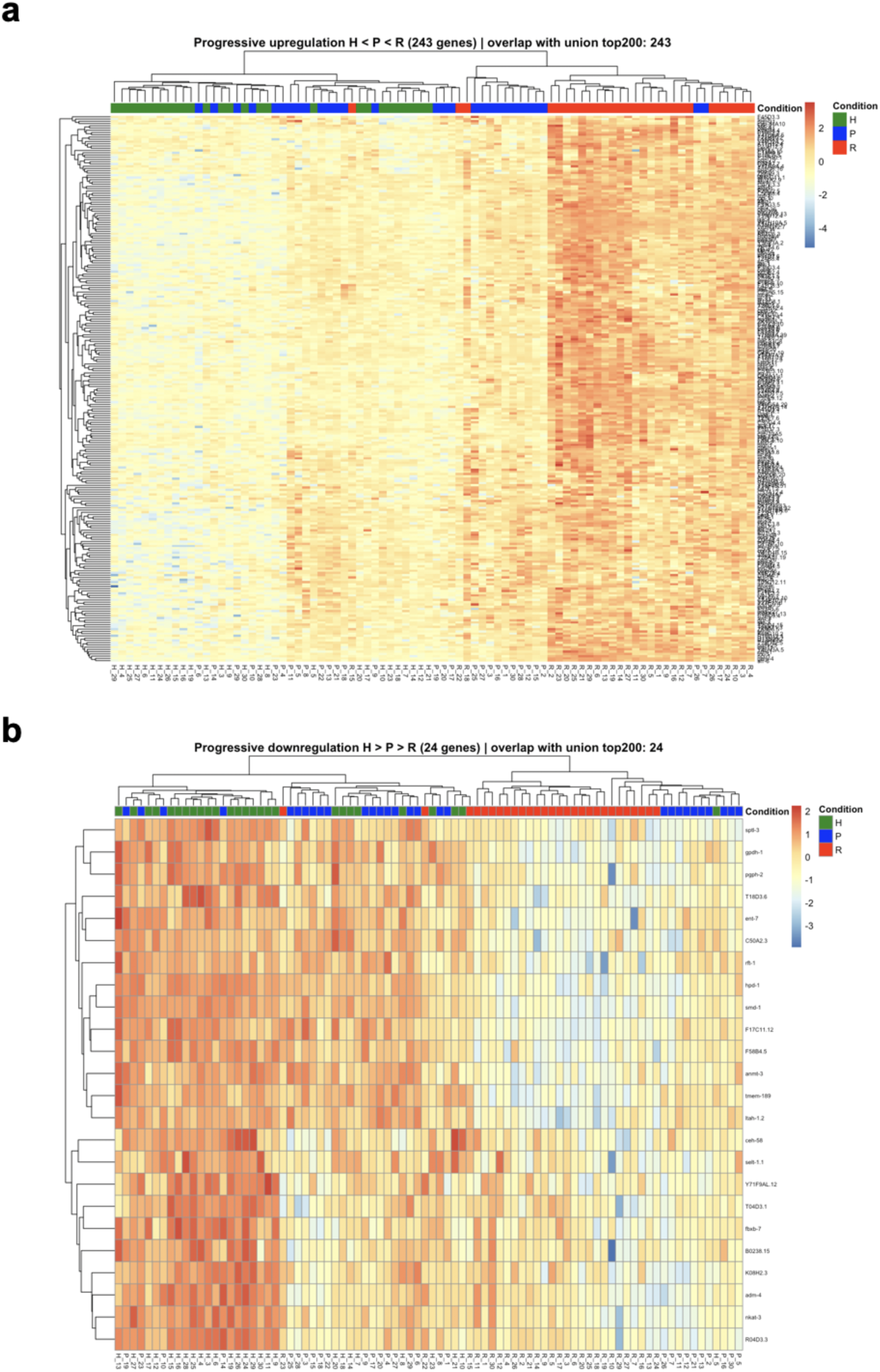
Single Worm somatic Top DE genes correlated with Rup and Pvl phenotypes. **a**) Heatmap of 243 genes showing progressive upregulation across phenotypic severity (mean H < mean P < mean R) **b**) Heatmap of the 24 somatic genes showing progressive downregulation across phenotypic severity (mean H > mean P > mean R). For **a**) and **b**) genes were selected from the union top 354 differentially expressed somatic genes in Rup and Pvl (DESeq2, padj < 0.05). Expression values are z-scored log normalized counts. Columns represent individual animals colored by phenotypic class; rows represent genes clustered by hierarchical clustering.

